# Prebiotic cationic amino acids support formation of an ancient protein fold

**DOI:** 10.64898/2026.07.28.741362

**Authors:** Sota Yagi, Aditya K. Padhi, Khushboo Bhagat, Kam Y. J. Zhang, Shunsuke Tagami

## Abstract

The evolutionary basis for the selection of the standard proteinogenic amino acids remains elusive, particularly for the long cationic amino acids, lysine and arginine, which were likely scarce in the prebiotic environment. In contrast, shorter cationic amino acids, such as ornithine (Orn), and 2,4-diaminobutyric acid (Dab), are thought to have been more abundant. Here, we investigated whether these prebiotic cationic amino acids can support protein tertiary structure formation, using computational protein design, biophysical and crystallographic analyses, and molecular dynamics (MD) simulations. We designed sequences for an ancient protein fold, the double-Ψ β-barrel (DPBB), with ornithine and plausible prebiotic amino acid sets. Although all the designed sequences were unfolded under standard dilute aqueous conditions, one Orn-containing variant folded in highly concentrated conditions. Crystallographic analysis revealed that both this peptide and its Dab-substituted derivative adopted the double-Z β-barrel (DZBB) fold, a likely evolutionary intermediate between extant β-barrel folds. Therefore, Orn and Dab might have supported the foldability of primitive proteins before the incorporation of lysine and arginine into the genetic code.

## Main

Life synthesizes proteins from twenty canonical amino acids designated by the genetic code, yet the rationale behind the evolutionary selection of these amino acids remains a fundamental mystery. It is generally believed that life evolved metabolic pathways to synthesize amino acids for protein production. However, in primitive organisms with immature metabolic systems, essential nutrients, including amino acids, were likely acquired from the surrounding environment^1–3^. Previous studies on prebiotic chemical evolution, represented by the volcanic spark discharge experiments, have demonstrated that several types of amino acids can be synthesized from inorganic compounds^4–7^. Moreover, given the identification of a wide variety of amino acids in carbonaceous meteorites, it is possible that these amino acids were supplied from extraterrestrial sources^8–10^. Although approximately half of the proteinogenic amino acids have been identified in these studies, some canonical amino acids remain undetected.

The modern genetic system encodes three cationic amino acids with relatively complex side chains: lysine, arginine, and histidine. These amino acids are critical for maintaining protein functionality, particularly by stabilizing tertiary structures and facilitating molecular interactions through electrostatic interactions. However, the prebiotic synthesis of Lys, Arg, and His remains highly challenging^11^, and these amino acids have not been detected in carbonaceous meteorites to date. Thus, their limited availability in the primordial Earth is particularly perplexing in the field of protein evolution^12,13^. In contrast, cationic amino acids with shorter side chains, such as ornithine (Orn), 2,4-diaminobutyric acid (Dab), and 2,3-diaminopropionic acid (Dap), have been synthesized by prebiotic chemistry and identified in meteorites (Fig. 1A)^9,14–16^. Although the possibility that lysine and arginine existed in ancient environments cannot be ruled out, it is more likely that such noncanonical cationic amino acids were more abundant on early Earth. Recently, it was reported that such non-canonical basic amino acids, in particular Orn and Dab, can also promote prebiotic energy-related chemistry, presumably by assisting proton delivery to the mineral surface^17^.

**Figure 1.**
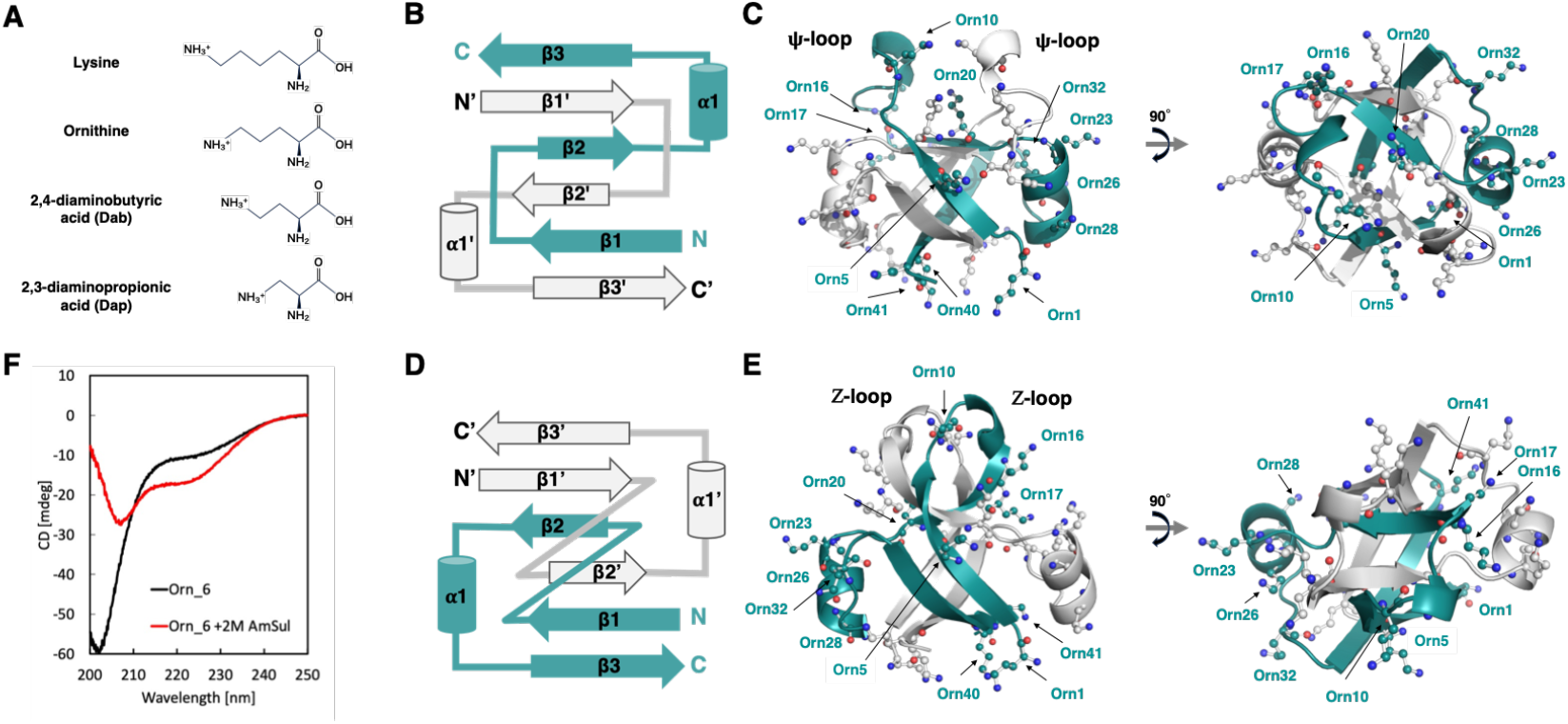
Design and structure of the ornithine peptide. (A) Chemical structures of lysine, ornithine, 2,4-diaminobutyric acid (Dab), and 2,3-diaminopropionic acid (Dap). (B) Topology of the homo-dimeric DPBB fold. (C) Cartoon representation of the lowest Rosetta score design (Orn_6) with the DPBB fold, illustrating the distribution of ornithine (Orn) residues. (D) Topology of the homo-dimeric DZBB fold. (E) Crystal structure of Orn_6, which adopts the DZBB fold. (F) Circular dichroism spectra of 500 µM Orn_6. Black and red lines show the spectra of the peptide in 50 mM phosphate, pH 6.0, and 150 mM NaCl, in the absence/presence of 2 M ammonium sulfate, respectively.

Interestingly, the modern bacterial lysyl-tRNA synthetase can charge not only lysine but also ornithine onto tRNA^Lys18^. Additionally, lysyl-tRNA synthase reportedly activates ornithine and Dab by conjugating AMP, but the pre-transfer editing system of the enzyme subsequently hydrolyzes the resulting complexes to avoid their incorporation into nascent peptides^19,20^. Given the scarcity of proteinogenic cationic amino acids and the relative abundance of ornithine, Dab, and Dap in prebiotic contexts, it is conceivable that these simple, positively charged amino acids could have been utilized in early protein synthesis.

To investigate this ancient scenario, peptides composed of non-canonical cationic residues were synthesized. Longo et al. demonstrated that ancient helix-loop-helix peptides with ornithine residues were unfolded but formed phase-separated droplets when mixed with a nucleic acid polymer^21^. Makarov et al. evaluated the solubility and secondary structure propensities of Dab in synthetic combinatorial peptide libraries and found that the addition of Dab to a short peptide sequence reduces its secondary structure potential^22^. Similarly, the helical propensities and the strengths of salt-bridges in secondary structures for ornithine, Dab, Dap, and lysine residues were also assessed^23–25^. These studies have provided valuable insights into the physicochemical properties of peptides containing non-canonical cationic amino acids. Building on these achievements, further investigation into their capability of forming well-defined protein folds will provide deeper insight into early protein evolution. As β-proteins, such as small β-barrel proteins, are essential components in the gene expression system, understanding their foldability with non-canonical cationic amino acids is crucial for elucidating protein evolution in the early life stage.

RNA polymerase is one of the indispensable enzymes in the gene expression system, and its catalytic core is composed of the small β-barrel fold called the double-Ψ β-barrel (DPBB). We previously demonstrated that the DPBB fold can be constructed by a homo-dimeric 43aa peptide with only seven amino acid types (Ala, Val, Gly, Asp, Glu, Lys, and Arg)^26^. This simplified sequence LysArg_0, previously described as mk2h_ΔMILPYS, was unfolded under typical buffer conditions but formed crystals in the presence of malonate or malic acid, adopting the homo-dimeric DPBB structure. Furthermore, LysArg_0 also crystallized with a high concentration of sulfate, and adopted a different fold named the double-Z β-barrel (DZBB)^27^. The fold conversion experiment suggested that this DZBB fold might be a missing-link fold connecting diverse β-barrel proteins.

In this study, we investigated how non-canonical cationic amino acids, such as ornithine, Dab, and Dap, affect the foldability of ancient β-barrel proteins. To this end, we designed and synthesized minimal DPBB/DZBB peptides incorporating these residues and examined their structural-dynamic properties by experimental and computational analyses. Our findings support the possibility that ancient proteins with shorter cationic amino acids existed as evolutionary intermediates during the early evolution of life.

## Result & Discussion

### Design and characterization of ornithine peptides

The simplified DPBB/DZBB peptide LysArg_0 was further engineered by substituting twelve positively charged residues (seven lysines and five arginines) with ornithine. The remaining amino acids were computationally optimized to maintain the homo-dimeric DPBB structure (Figs. 1B and C, Fig. S1). During this computational design process, the amino acid sets were constrained to either (subset 1) the five amino acid types (Ala, Val, Gly, Asp, and Glu) encoded by GNN codons in the standard genetic code or (subset 2) the ten amino acid types presumed to be prebiotic, resulting in seven designs in total (Orn_1–7) (Table S1 and Fig. S1). The details of the sequence designs are provided in the supplementary information (Supplemental information).

The seven designed Orn-peptides were dissolved in phosphate buffer (50 mM potassium phosphate, pH 6.0, 150 mM NaCl) and subsequently analyzed for their biophysical properties. In the size exclusion chromatography (SEC) experiment using a Superdex™ 75 Increase 10/300 column, all ornithine-substituted peptides eluted at approximately 13–14 mL, indicating unfolded or aggregated states (Fig. S2). Similarly, in the circular dichroism (CD) analysis, all peptides exhibited characteristics of random coils (Fig. S3).

Since LysArg_0 formed diffraction-quality crystals, even though it was unfolded in the SEC and CD experiments^6^, crystallization screening of all ornithine-substituted peptides was carried out using three screening kits (288 buffer conditions in total). Only Orn_6, composed of ten prebiotic amino acids including twelve ornithine residues, crystallized under conditions with 40% (v/v) PEG 400, 100 mM Tris-HCl (pH 8.5), and 200 mM lithium sulfate, and its crystal structure was determined at 1.8 Å resolution (Table S2). Unexpectedly, Orn_6 adopted the DZBB conformation (Fig. 1D and E), rather than the DPBB structure used as the model during the computational design process (Fig. 1B and C). A similar unanticipated conformational change between DPBB and DZBB was observed in a previous experiment, where LysArg_0 adopted either the DPBB or DZBB structure under different crystallization conditions^27^. Although the structural basis underlying the observed divergence of Orn_6 from the designed DPBB fold remains unclear, unexpected folding pathways or kinetic traps, which could not be accounted for during the design process, may have led to the formation of the DZBB structure.

To test whether the folding of Orn_6 is induced by the high peptide and salt concentrations used during crystallization, we examined its CD spectra under concentrated conditions. At a peptide concentration of 500 µM, Orn_6 remained in a random coil state in the absence of salt. By contrast, it exhibited non-random coil features in the presence of 2 M ammonium sulfate (Fig. 1F). These observations suggest that high peptide concentrations in combination with sulfate promote folding, possibly by increasing intermolecular encounters, and thereby facilitating formation and stabilization of the homo-dimeric structure. Consistent with this interpretation, folding of the parent peptide LysArg_0 was also enhanced under high sulfate conditions^27^. Collectively, these results indicate that Orn_6 is capable of adopting a folded conformation not only in the crystalline state but also in highly concentrated aqueous conditions.

In the asymmetric unit of the Orn_6 crystal structure, two peptide chains form the homo-dimeric DZBB fold (Fig. 1E). When comparing the two chains, the fundamental structures are similar, but the directions of α1 in both chains differ slightly and are accompanied by the stretching of the connecting loop between α1 and ?3 (Fig. S5). This difference implies that the helical region and the loop are more flexible in comparison with the core barrel. We previously reported that the peptide without the α1 sequence maintained its stable DZBB fold, which might serve as an evolutionary intermediate to a different β-barrel fold, the OB-fold^27^. Therefore, we constructed an Orn_6 mutant (Orn_6_woH) by trimming the six residues corresponding to the α1 region in Orn_6 (Fig. S6A). Orn_6_woH was unfolded under the standard buffer conditions, but like Orn_6, it formed crystals and adopted the DZBB fold (Fig. S6B and C). Notably, it also exhibited a non-random coil CD spectrum at high peptide concentration in the presence of 2M ammonium sulfate (Fig. S4A). These results imply that ornithine-containing peptides could access an intermediate structural state that may have bridged the DZBB and OB folds during protein fold evolution.

### Investigation of peptides with varied side chain lengths

To examine the foldability of peptides with other prebiotic cationic residues, we further engineered Orn_6 by replacing its twelve ornithine residues with Dab or Dap (Table S1). The resultant mutants are termed Dab_6 and Dap_6, respectively. The biophysical analysis revealed that both peptides are unfolded under the standard buffer condition and remained unfolded even at high peptide concentration in the presence of 2M ammonium sulfate. (Fig. S7A and B, S4B and C). In the crystallization screening, only Dab_6 formed crystals in 2400 mM sodium malonate dibasic. We determined its crystal structure and confirmed that it adopts a homo-dimeric DZBB structure, similar to Orn_6 (Fig. 2A) (Table S2). As observed in the Orn_6 structure, the angle of the helix region in chain B slightly differs from that in chain A (Fig. S7C). Therefore, the short peptide including Dab could fold into a homo-dimeric ?-barrel structure similar to that of the ornithine peptide.

**Figure 2.**
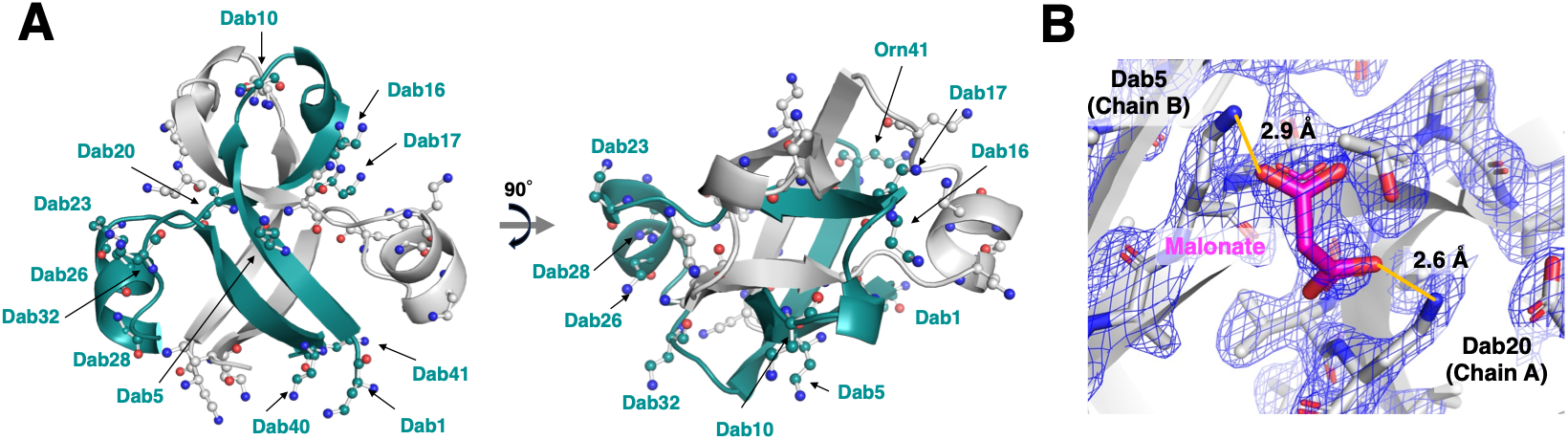
Crystal structure of the Dab_6 homo-dimer. (A) All Dab residues are shown as stick and ball models, and only Dab residues in chain A (cyan) are labeled. (B) Malonate is shown as magenta sticks and interacts with amino groups from Dab5 and Dab20 side chains. The composite omit electron density map for malonate is shown as a blue mesh contoured at 1.5σ.

In contrast to Orn_6, folding of Dab_6 was not detected under high peptide and ammonium sulfate concentrations based on CD measurements (Fig. S4B). This observation suggests that sulfate is insufficient to promote the folding and stabilization of Dab_6 in solution. Instead, the crystallization condition containing malonate may play a specific role in promoting its folding. Malonate, a kosmotropic dianion with two carboxylate groups and a larger molecular framework than sulfate, might facilitate stabilization through multivalent electrostatic interactions. Although CD measurements in such conditions could not be performed due to strong UV absorption by malonate, electron density corresponding to malonate molecules is clearly observed in the crystal structure of Dab_6. Notably, one malonate bridges two Dab side chains from adjacent β-strands, forming a cross-strand interaction (Fig. 2B).

We also performed molecular dynamics (MD) simulations to evaluate the structural stabilities of all three systems, Orn_6, Dab_6, and Dap_6, using the crystal structures of Orn_6 and Dab_6, and the modeled structure of Dap_6 as the starting structures (Fig. 3). The normalized root mean square deviation (RMSD) and radius of gyration (Rg) plots showed that both Orn_6 and Dab_6 maintained DZBB folds throughout the simulation (Fig. 3A and B and Fig. S8A and B). In addition, the free energy landscape analysis based on principal component analysis (PCA) further revealed that they exhibited compact energy basins with low trace values of 1.67 nm^2^ and 1.90 nm^2^, respectively, indicating restricted motion and stable folding consistent with the maintenance of the DZBB fold (Fig. 3C). In contrast, Dap_6 exhibited higher RMSD and Rg values, with a higher PCA-derived trace value (40.85 nm^2^), accompanied by a reduced number of hydrogen bonds and the loss of native secondary structure elements (Fig. 3B–F). These findings suggest that, unlike Orn and Dab, Dap is incapable of stabilizing the DZBB structure. This observation implies that the side-chain length of cationic residues may critically influence protein foldability.

**Figure 3.**
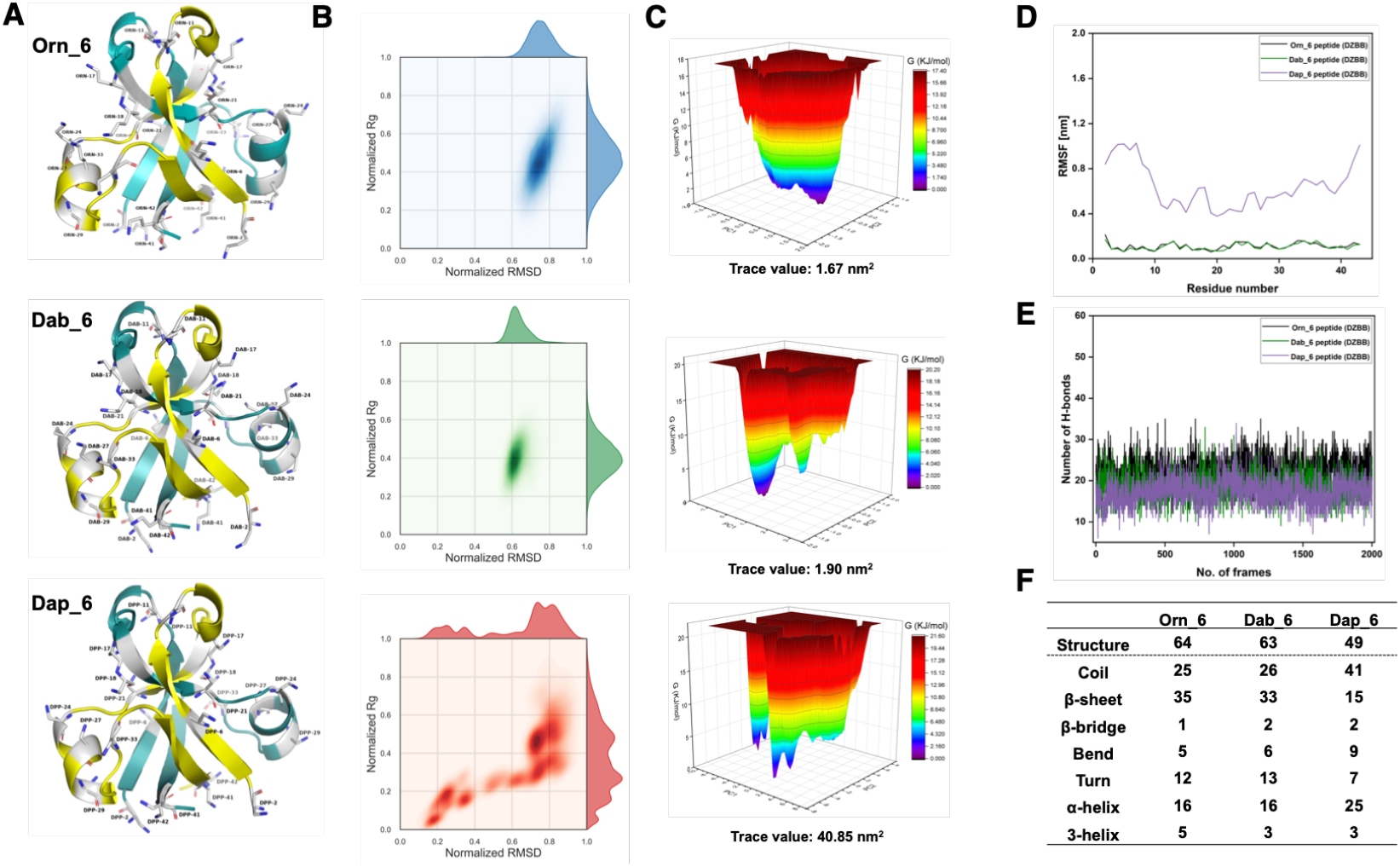
Molecular dynamics simulations of Orn_6, Dab_6, and Dap_6. (A) Representative structures with the DZBB form used for the MD simulations. (B) Normalized RMSD vs. Rg plots. (C) Free energy landscapes constructed from the dominant principal components. The first two principal components capture the essential collective motions, and the corresponding trace values reflect differences in conformational sampling. Panels (A) to (C) show representative figures for Orn_6, Dab_6, and Dap_6, respectively. (D) Root mean square fluctuation (RMSF) profiles. (E) Intramolecular hydrogen bonds across 20,000 frames. (F) Secondary structure elements (%). Structure = α-helix + β-sheet + β-bridge + Turn.

Subsequently, we redesigned the Orn_6 peptide by substituting all ornithine residues with lysine residues, producing the Lys_6 peptide (Table S1 and S3). In the SEC experiment, Lys_6 exhibited a single peak corresponding to a dimer (Fig. 4A). The CD experiment revealed that Lys_6 formed a non-random-coil structure and denatured cooperatively in a two-state manner, even in the standard dilute condition (Fig. 4B). Its melting temperature was calculated as 91°C, indicating its high thermostability (Fig. S9). The one-dimensional proton NMR spectrum of Lys_6 exhibited well-dispersed chemical shifts, characteristic of a well-defined tertiary structure (Fig. 4C). Therefore, unlike Orn_6 and Dab_6, Lys_6 formed a stable structure with high thermostability under standard buffer conditions. However, we could not determine whether its topology is DPBB or DZBB, as we could not obtain its crystal, possibly because the longer flexible side chains on the protein surface hindered its crystallization.

**Figure 4.**
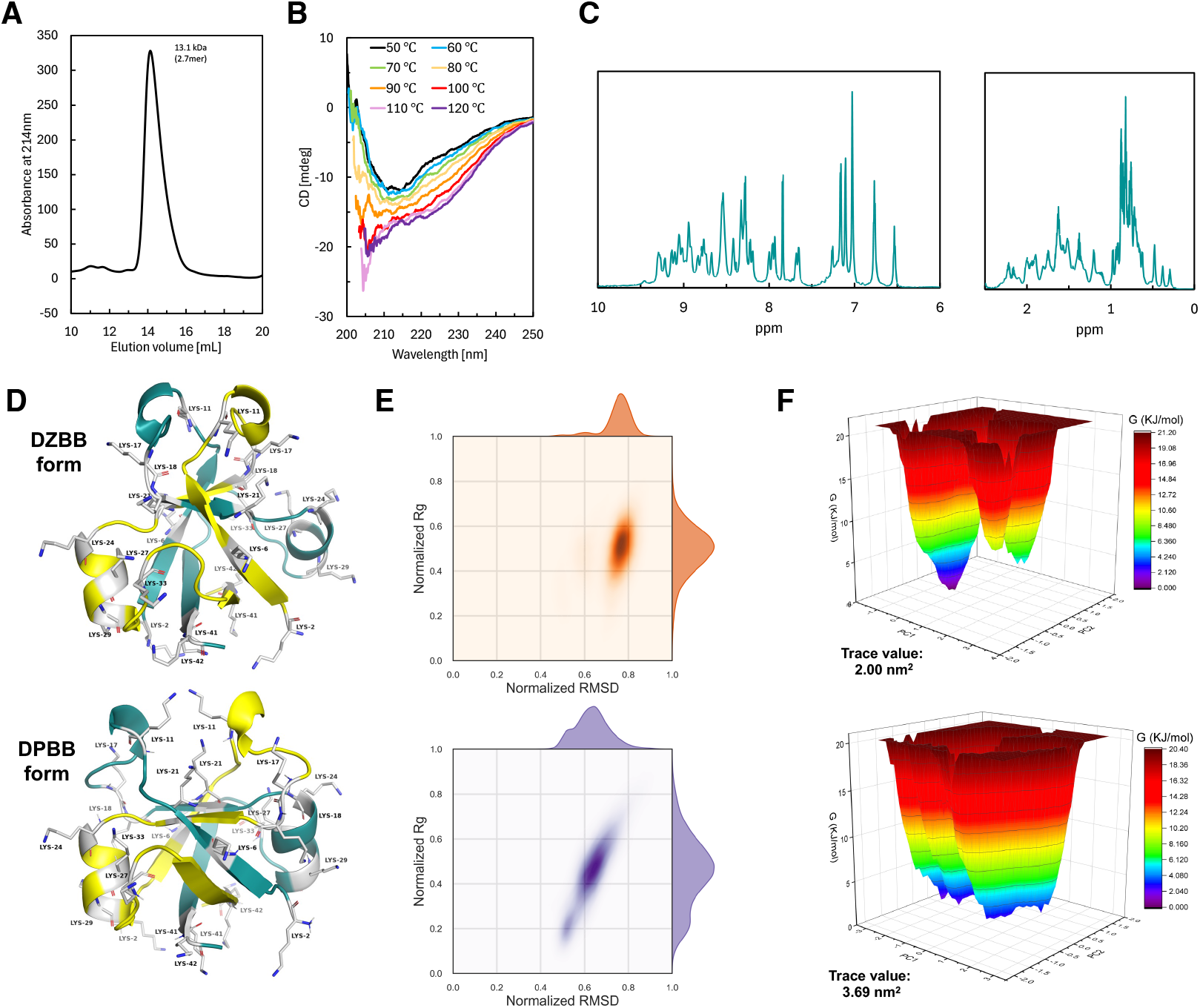
Physicochemical characterization of Lys_6. (A) Size-exclusion chromatogram. The apparent molecular mass was calculated to be 13.1 kDa. (B) CD spectra at different temperatures. (C) One-dimensional proton NMR spectrum showing peaks for amide protons (left panel) and the aliphatic region (right panel). (D) Representative structures with the DZBB and DPBB forms for the simulations. Panels (E) and (F) shows the Normalized RMSD vs. Rg plots and Free energy landscape analysis profiles of Lys_6 in the DZBB- and DPBB-forms, respectively.

To estimate the structural stabilities of Lys_6 in the DPBB and DZBB forms, we performed further MD simulations (Fig. 4D and E; Fig. S8). Both conformational states exhibited comparable stabilities with low RMSD, Rg, and PCA-derived trace values and preserved secondary structure elements throughout the trajectories (Fig. 4E, F, and Fig. S8C). Taken together with the experimental results, these findings suggest that Lys_6 possesses the intrinsic ability to form stable β-barrel structures, although its precise topological arrangement (DPBB or/and DZBB) remains uncertain.

### Binding properties to nucleic acid polymers

Cationic residues induce interactions with negatively charged nucleic acid polymers through electrostatic interactions. Thus, we assessed whether the peptides designed in this study could bind to nucleic acids in a dilute condition (5 µM peptides), using an electrophoretic mobility shift assay (EMSA).

Among the seven ornithine peptides, only Orn_3 slightly slowed down the migrations of dsDNA, and ssDNA, and dsRNA (Fig. 5A)(Table S3). We also investigated the binding ability of Dab_6, Dap_6, and Lys_6 to nucleic acids and found that Lys_6 shifted the mobilities of dsDNA, ssDNA, and partially double-stranded RNA significantly (Fig. 5B). In particular, the dsDNA formed a large, aggregation-like complex with Lys_6 and remained in the loading well of the gel. Given that the nucleic acid affinities for oligolysine and oligoornithine peptides are generally comparable^28^, it is reasonable to infer that the DNA-binding ability of Lys_6 arises from its folded conformation in the dilute condition.

**Figure 5.**
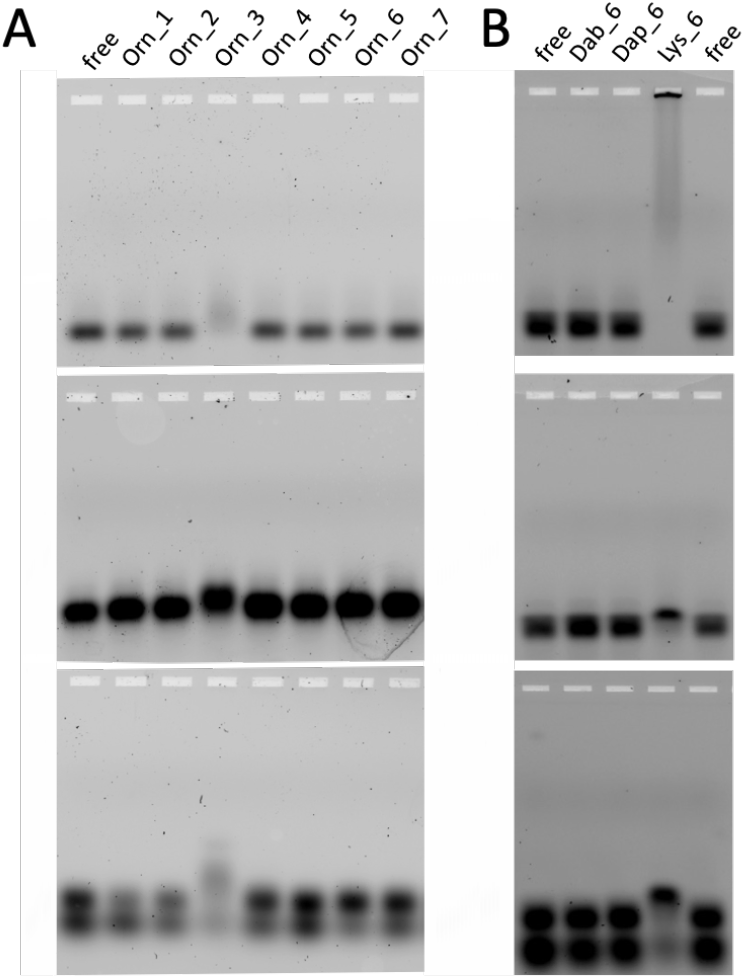
Characterization of the nucleic acid binding properties of the peptides. (A) Orn_1– 7 (5 µM) or (B) Dab_6, Dap_6, or Lys_6 (5 µM) was mixed with FAM-labeled double-stranded DNA (top), single-stranded DNA (middle), or partially double-stranded RNA (bottom). The resulting peptide–nucleic acid complexes were analyzed by an electrophoretic mobility shift assay (EMSA) on a 2% agarose gel.

## Conclusion

This study demonstrated the construction of the ancient β-barrel structure DZBB, using the simple and non-canonical cationic amino acids Orn and Dab and nine other amino acids that would have been abundant in a prebiotic environment, instead of using Lys and Arg in the modern genetic code (Figs. 1 and 2). In contrast, the shortest cationic residue tested, Dap, could not support the formation of the ancient β-barrel structure (Fig. 3).

Previous studies indicated the chemical limitations of Orn and Dab due to their pH- and activation-dependent susceptibility to intramolecular lactamization. In their free or peptide-incorporated forms, Orn and Dab are relatively stable under neutral aqueous conditions but tend to cyclize under alkaline conditions, where the deprotonated side chain amine attacks the backbone carboxylic group^29–31^. In contrast, when these non-proteogenic amino acids are activated for polymerization–for example, as adenylated or nitrile intermediates–they become highly prone to rapid intramolecular lactamization even at neutral pH, which hinders further peptide polymerization^13,32–34^. Nevertheless, once these residues were incorporated, our results indicate that they could still give rise to well-defined tertiary structures. This implies that Orn and Dab may have acted as ancient proteinogenic residues that enabled the formation of primitive β-barrel folds—such as DZBB—at a time when Lys was scarce or absent in prebiotic or early biosynthetic environments. Although Orn_6 was originally designed to adopt the DPBB fold, Orn_6 and Dab_6 instead formed the DZBB fold, which is not observed in extant proteins but has been proposed as a missing-link architecture capable of structural interconversion with other β-barrel folds, including RIFT and OB folds (Fig. S10). In this context, Orn and Dab may have contributed to maintaining structural robustness and evolutionary continuity in primitive proteins prior to the full establishment of coded protein synthesis.

Although Orn- and Dab-containing peptides appeared unfolded under standard dilute aqueous conditions (Fig. S3), their folding behavior was strongly dependent on environmental conditions. In particular, Orn_6 and Orn_6_woH exhibited non-random coil CD features under a high peptide concentration in the presence of 2M ammonium sulfate (Fig. 1F and S4A). Likewise, Dab_6 adopted the DZBB fold with malonates in the crystalline state (Fig. 2B). Intracellular environments are highly crowded and contain various salts and metabolites that can affect protein folding and stability^35,36^, unlike the simple buffer conditions. Similarly, under prebiotic settings such as a dry-wet cycle, local peptide and salt concentrations could have become extremely high^37–39^. Thus, the ability of Orn- or Dab-containing peptides to fold in concentrated environments suggests that their structural potential may have been realized under physiologically or prebiotically relevant conditions, rather than under dilute conditions used in typical biophysical experiments.

By contrast, the substitution with Lys markedly enhanced the structural stability even under dilute aqueous conditions (Fig. 4), and the Lys-containing peptide exhibited affinities toward dsDNA, ssDNA, and partially double-stranded RNA (Fig. 5). These results suggest that the recruitment of Lys into the genetic code was not only due to its greater chemical stability than Orn and Dab but also because its longer hydrocarbon side chain enabled the formation of more favorable salt-bridges with negatively charged residues^25,40^ and contributed additional hydrophobic interactions^25,41^, thereby allowing proteins to function under diverse environmental conditions as observed today. Thus, the transition from Orn/Dab to Lys may represent a critical evolutionary step from ancient proteins with low synthetic costs to more robust and functional molecular machines.

## Acknowledgments

This work is based on experiments performed at KEK (project number: 2022G005) and SPring-8. The authors are grateful to the beamline staff scientists at KEK and SPring-8. We thank Toshiaki Hosaka and Kentaro Ihara for assistance with the X-ray diffraction experiments. We also thank Toshio Nagashima for assistance with the NMR experiment. S.Y. and S.T. were supported by JSPS (22H01346 and 24K07206). S.T. was also supported by the Astrobiology Center Program of National Institutes of Natural Sciences (AB0713). S.Y. was also supported by the JGC Saneyoshi Research Grant Program (2419). K.B. acknowledges the Ministry of Human Resource Development (MHRD), Government of India, and IIT (BHU) for her research fellowship. A.K.P. is grateful for financial support received from the Science & Engineering Research Board (SERB), Government of India’s Project No. SRG/2023/000167. Further, the computing resources of PARAM Shivay Facility under the National Supercomputing Mission, Government of India, at the IIT (BHU), Varanasi, are gratefully acknowledged.

## Notes

### Competing Interest Statement

The authors have declared no competing interest.

